# An Automated Workflow for Segmenting Single Adult Cardiac Cells from Large-Volume Serial Block-Face Scanning Electron Microscopy Data

**DOI:** 10.1101/242701

**Authors:** Akter Hussain, Shouryadipta Ghosh, Siavash Beikoghli Kalkhoran, Derek J. Hausenloy, Eric Hanssen, Vijay Rajagopal

**Author notes:** corresponding author; Level 4, Building 170, Department of Biomedical Engineering, The University of Melbourne, Parkville, VIC 3010.

## Abstract

This paper presents a new algorithm to automatically segment the myofibrils, mitochondria and nuclei within single adult cardiac cells that are part of a large serial-block-face scanning electron microscopy (SBF-SEM) dataset. The algorithm only requires a set of manually drawn contours that roughly demarcate the cell boundary at routine slice intervals (every 50^th^, for example). The algorithm correctly classified pixels within the single cell with 97% accuracy when compared to manual segmentations. One entire cell and the partial volumes of two cells were segmented. Analysis of segmentations within these cells showed that myofibrils and mitochondria occupied 47.5% and 51.6% on average respectively, while the nuclei occupy 0.7% of the cell for which the entire volume was captured in the SBF-SEM dataset. Mitochondria clustering increased at the periphery of the nucleus region and branching points of the cardiac cell. The segmentations also showed high area fraction of mitochondria (up to 70% of the 2D image slice) in the sub-sarcolemmal region, whilst it was closer to 50% in the intermyofibrillar space. We finally demonstrate that our segmentations can be turned into 3D finite element meshes for cardiac cell computational physiology studies. We offer our large dataset and MATLAB implementation of the algorithm for research use at www.github.com/CellSMB/sbfsem-cardiac-cell-segmenter/. We anticipate that this timely tool will be of use to cardiac computational and experimental physiologists alike who study cardiac ultrastructure and its role in heart function.

## INTRODUCTION

Serial block-face scanning electron microscopy imaging (SBF-SEM) is a technology that has enabled full three-dimensional analysis of large tissue blocks at intermediate resolution (10–50nm) bridging the gap between high resolution electron tomography and optical microscopy serial sections. Studies by Pinali et al^1–3^ provided new views of the intricate sarcoplasmic reticular network in cardiac cells. Glancy and collaborators^4^ used serial block-face image data to examine the 3D arrangement of mitochondria in skeletal muscle tissue blocks, while Holzem et al^5^ used 3D SBF-SEM images to explore differences in cardiac ultrastructure between control and heart failure patients. The three-dimensional and high throughput nature of SBF-SEM imaging make the technique an invaluable resource for any structure-based analysis of subcellular mechanisms in health and disease.

The amount of data generated leads to the necessary but time consuming task of segmenting the ultrastructural details from these very large datasets. Indeed, the studies cited above refer to a combination of manual, threshold-based and machine-learning-based segmentation to extract specific components of their cellular datasets. Currently used segmentation tools (which we also use) include Fiji^6^, IMOD^7^ and ilastik^8^, to name a few. However, a user still has to overcome the challenge of identifying the appropriate combination of algorithms and parameters to segment a particular SBF-SEM cardiomyocyte dataset using these tools. We present a new method for automated segmentation of cardiac ultrastructure from SBF-SEM imaging data that requires minimal user interaction. Furthermore, we have developed our algorithm with segmentation of individual cells in mind as opposed to an indiscriminate segmentation of a tissue region, which is the norm with all other existing segmentation tools. This enables single-cell based analyses as well as analyses of emergent properties of collections of cells.

We have used this algorithm to segment the nuclei, myofibrils and mitochondria of three heart cells contained within two SBF-SEM-acquired datasets that were collected at the University of Melbourne, Australia. One of these datasets spans a volume of 215 μm × 46 μm × 60 μm at 50 nm isotropic voxel spacing is, as far as we are aware, the largest dataset of a single cardiac cell acquired using SBF-SEM. We demonstrate the improved performance of our algorithm for cardiac cell segmentation relative to algorithms used by IMOD and Fiji. We also provide information on the sensitivity of the algorithm to images with different contrasts by also testing the algorithm on two additional datasets: one dataset was acquired at University College London, independent of this study on a different block face imaging system; and another is a recently published and freely available dataset^9^.

We provide the MATLAB implementation of the algorithm as well as the segmented dataset of myofibrils, mitochondria and nuclei free to use at our GitHub repository at www.github.com/CellSMB/sbfsem-cardiac-cell-segmenter/. We anticipate that the segmented dataset will prove invaluable to mathematical modelers interested in studying cardiac biophysics in spatially detailed models of cardiac ultrastructure^10–12^. We believe that our MATLAB implemented algorithm will prove useful to many cardiac biophysicists who are interested in rapid extraction and analysis of ultrastructure content and organization without manual segmentation or prior knowledge of image processing techniques. We also encourage readers to implement the general principles of the algorithm in other existing tools such as IMOD or Fiji, which would provide many additional tools for further processing.

## METHODS

The following methods on tissue preparation and serial block-face imaging were used to collect the SBF-SEM data at the University of Melbourne and University College London.

### Tissue Sample Preparation

The tissue samples used for SBF-SEM imaging were prepared at the University of Auckland. All animal procedures followed guidelines approved by the University of Auckland Animal Ethics Committee (for animal procedures conducted in Auckland, Application Number R826).

Sixteen-week-old male Sprague-Dawley rats were euthanized and hearts were excised and quickly cannulated and connected to a Langendorff apparatus operated at 90 cm hydraulic pressure. The heart was perfused with Tyrode solution including 20 mM 2,3-Butanedione Monoxime for 2–3 minutes followed by perfusion with 2.5% glutaraldehyde, 2% paraformaldehyde, 50 mM CaCl_2_, in 0.15 M sodium cacodylate buffer.

Two tissue blocks were dissected from the left ventricular free wall and stored in the same fixative that had been precooled on ice for 2 hours. The fixative was subsequently replaced with 2% osmium tetroxide and .8% potassium ferrocyanide in 0.15 M sodium cacodylate and left overnight. The blocks were then stained with ice-cold 2% uranyl acetate for 60–120 minutes. The blocks were then washed of excess uranyl acetate and punched into 1.5 mm diameter samples. The samples were then progressively dehydrated with ethanol followed by transition to room temperature in acetone. The samples were then embedded in epoxy resin (Durcupan ACM resin from EM sciences).

We tested the robustness of our algorithm on a dataset from University College London that was collected independent of this study. The experiments from which this dataset was acquired were conducted in compliance with the Animal (Scientific Procedures) Act 1986 published by the UK Home Office. The details of the sample preparation are provided as follows: A harvested heart from a 10–14-week-old C57/BL6 mouse was cannulated, washed with Krebs-Henseleit buffer containing NaCl (118 mM), NaHCO_3_ (25 mM), d-Glucose (11 mM), KCl (4.7 mM), MgSO_4.7_H_2_O (1.22 mM), KH_2_PO_4_ (1.21 mM) and CaCl_2_.H_2_O (1.84 mM) and fixed (1% paraformaldehyde, 2% glutaraldehyde in 0.1M sodium cacodylate buffer). The tissue was then processed for SBF-SEM based on a different protocol to the one used at the University of Melbourne^13^.

### Serial Block Face Imaging

With the samples collected at the University of Melbourne, the orientation of the muscle fibers was found using microCT (GE nanotom microCT, TrACEES platform, The University of Melbourne) of the whole resin block. The block was then trimmed to a square block face of 1 mm^2^ and 300 μm deep with fibers running perpendicular to the future cutting face. The trimmed block was then silver glued on a SEM stub (Agar Scientific, AGG1092450). The sample was mounted on a Teneo VolumeScope (FEI, Hillsborough, USA) and imaged in low vacuum mode (50 mbar) at 3kV, 0.1nA current using a backscattered detector. The acquisition pixel size was set at 10 nm for x and y.

Example slices from the two tissue blocks, showing the 3 cells, are provided in Figure 1. For tissue block 1, one thousand and nineteen serial sections of 50 nm thickness were acquired. The sections were aligned using IMOD^7^. The data was binned to an isotropic voxel size of 50 × 50 × 50 nm. and rotated 90 degrees around the Y-axis The image stack for tissue block 2 was binned to the same isotropic voxel resolution and contains 506 sections with 1024x884 pixels in each section. Cell 1 contains the largest volume of heart cell structure information as the dataset for tissue block 1 captures the entire cell cross section and the entire length of the cell. The datasets only capture part of the volume of cells 2 and 3.

**Figure 1:**
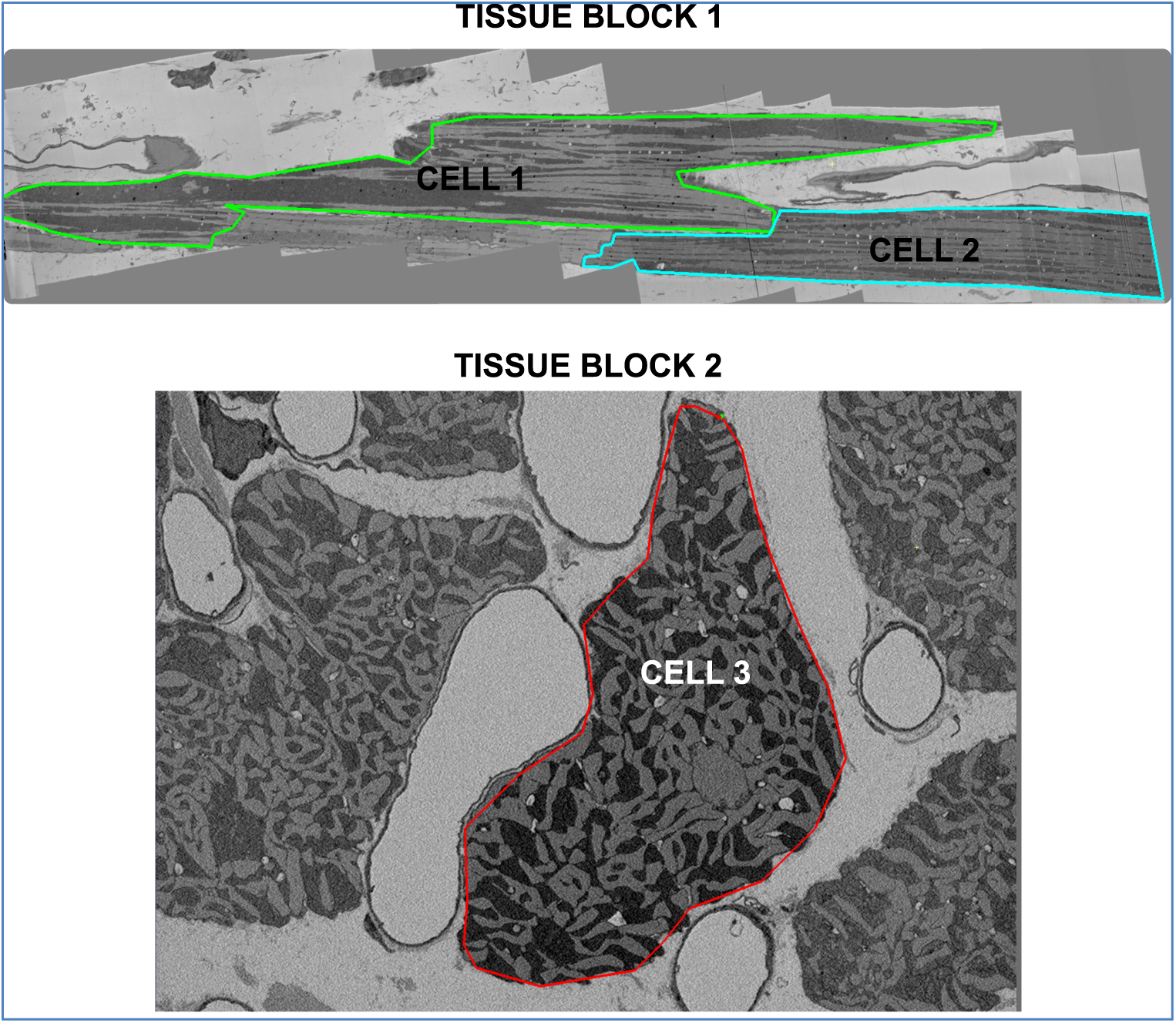
Cells used for the study. A total of 3 cells across two tissue blocks, as illustrated in this figure, were acquired using SBF-SEM.

The processed tissue that was collected at University College London was mounted on a pin and cut to 1 × 1 mm dimensions covering cardiomyocytes in their longitudinal orientation. After silver coating, the sample was placed in the Gatan 3 view within Zeiss Sigma variable pressure field emission SEM. Finally, samples were serially sectioned with the resolution of 100nm in Z and 32 × 32 nm in X and Y. Captured images were aligned using Gatan Digital Micrograph software.

### Single Cell Segmentation Algorithm

The 5 core steps of the algorithm are stated and described below and outlined in Figure 1.

#### Step 1, Cell boundary estimation

The algorithm begins with a set of manually demarcated cell boundaries from the SBF-SEM image stack (Figure 2A). We manually segmented every 10^th^ slice of the 591 slices that make up the cell volume using IMOD ^7^. Figure 2B shows a close-up view of one of the manually drawn contours of the cell boundary and highlights a strength of the proposed algorithm. As shown, the contours do not have to precisely follow the cell boundary or intercalated disc. The algorithm uses these manual segmentations as initial guesses at the cell boundary in neighboring image slices. After refinement of this initial guess (see step 4) and successful segmentation of a slice the segmented boundary is propagated to the next slice as the initial guess.

**Figure 2:**
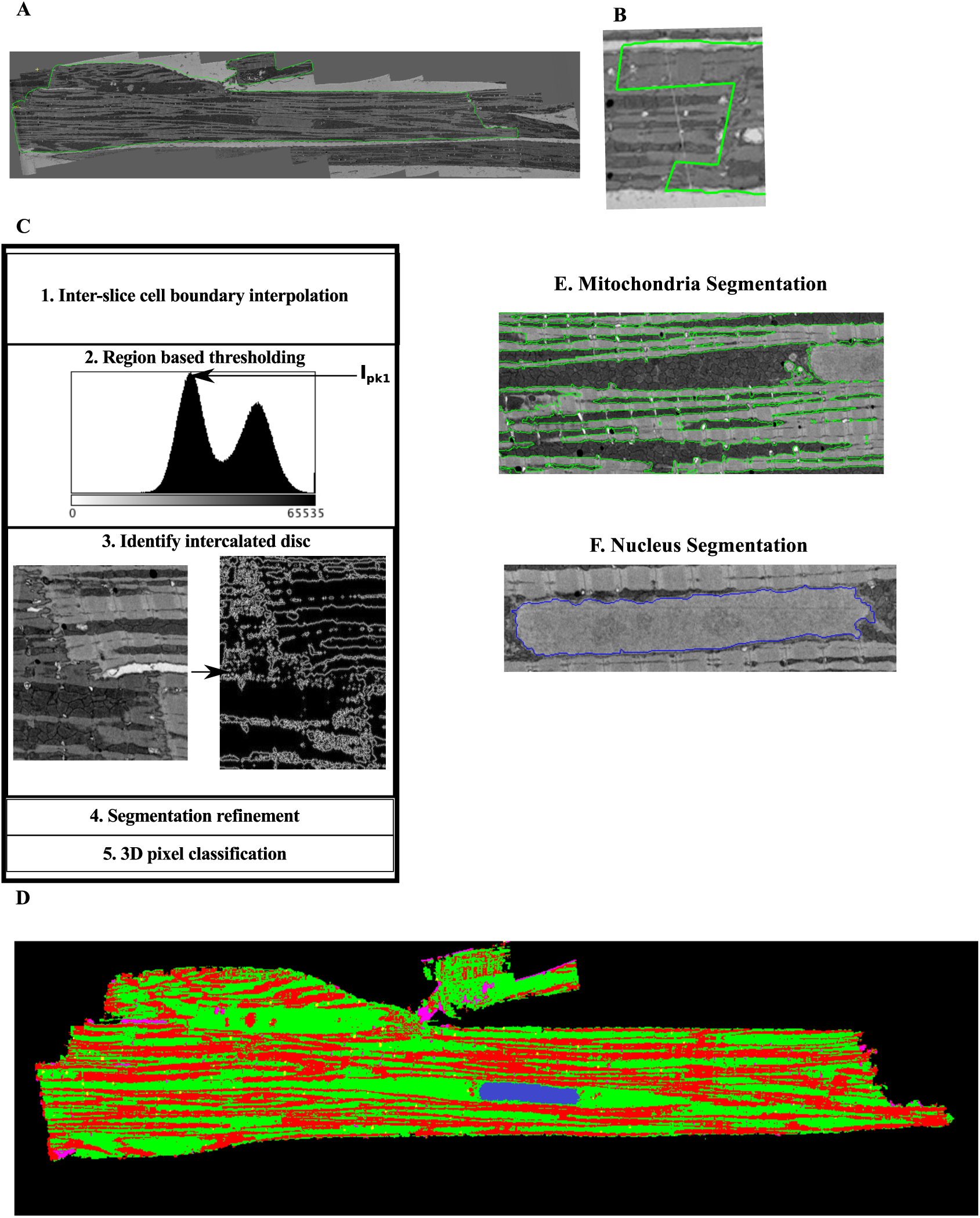
Workflow of volumescope segmentation. (A) An image slice showing manually drawn cell boundary contour in green; (B) a close-up view of a manually drawn contour near an **intercalated disc** to highlight the roughness of the contour; (C) steps towards a segmented image in (D). C(2) shows a typical histogram of intensities within an image. The two peaks correspond to mitochondrial and myofibrillar intensities; C(3) shows that calculating the intensity gradient near intercalated discs generates a dense region of non-zero gradient pixels; (D) segmented myofibrils in red, mitochondria in green, nucleus in blue and the remaining unclassified pixels in pink; (E) The green lines depict the

#### Step 2, Region-based thresholding

We implemented an automated region-based threshold algorithm to segment the pixels identified within the demarcated region of interest. In this step, we calculate an adaptive threshold that splits the image into two major components: myofibrils and mitochondria.

All pixels within the demarcated contour are sorted in ascending intensity values and stored in an array, *I.* Another array containing the pixel frequency distribution for the range of intensities within the image is also constructed. As shown in Figure 2C, step 2, the intensity histogram shows two distinct peaks that we identified as mitochondrial and myofibrillar.

In order to split the image region into these two organelle groups, the first peak (associated with mitochondria), *I_pk_*_1_, is determined by identifying the intensity value within the first half of the intensity range that the maximum number of pixels hold. If *I_pk_*_1_ were to be set as the threshold, it would ignore pixels which are still mitochondrial but are above *I_pk_*_1_. To correct for this under-estimation the mitochondrial threshold, *TH_mito_* is computed as *I_pk_*_1_ + (*I_pk_*_1_ − *I_min_*) where *I_min_* is the minimum intensity value in the region of interest. (*I_pk_*_1_ − *I_min_*) is the range of intensities that are below the first peak that have also been included as mitochondrial. The corrected *TH_mito_* includes a similar range of pixels above *I_pk_*_1_ to correct for variations in mitochondrial intensity.

An adaptive threshold is then computed using the formula *TH_adapt_* = max(*TH_mito_*, *I_median_*) where ***I****_median_* is the median intensity value of the region of interest. This threshold is then applied on the entire image – including the pixles outside the current estimate cell region – to create a first segmentation of the mitochondria region, which have intensity values less than the adaptive threshold.

#### Step 3, Gradient of the region-thresholded binary image and intercalated disc identification

Intercalated discs are undulating and narrow while mitochondria are larger, and more elliptic in shape. Therefore, mitochondrial edges, marked by a high intensity gradient, are fewer pixels thick, while pixels of intercalated discs will be identifiable by the regional pixels that have a non-zero gradient within the vicinity of each other. This distinction enabled us to separate regions that were mitochondrial from intercalated discs.

Figure 2C, Step 3 shows the result of calculating the intensity gradient on the thresholded image in Step 2. The algorithm identifies the dense areas depicted in the image as possible intercalated discs. Returning to the output of step 2, all pixels corresponding to possible gaps of intercalated discs that are found in step 3 and are outside the current estimated cell region, are removed.

#### Step 4, Segmentation refinement

At this step of the algorithm, the estimated cell boundary and region-based threshold image of mitochondria are used to produce a refined segmentation result.

First, the pixels of the region-based thresholded binary image from step 3 are grouped into different connected regions. The image is then subjected to iterative morphological erosion and dilation. The size of the erosion/dilation window is increased by 2 pixels at the start of each erosion/dilation step. After an erosion, all pixels that are outside the region of interest are classified as outside the cell. Any binary connected region for which 70% or more of its area is within the region of interest is classified as within the cell. Subsequently, the image is dilated back by the same window size. This operation of erosion, classification of regions inside and outside the cell and subsequent dilation is repeated until all regions have been classified.

#### Step 5, 3D pixel classification

After all pixels in the current 2D image have been classified, we further refine the segmentation by steps 1–4 in the other two orthogonal planes of the collected volume data. A pixel is now only classified as mitochondrion if it is categorized as mitochondrial in at least two of the three orthogonal planes.

With each image slice divided into a mitochondria or myofibril region, the pixels contained within the nucleus is typically classified as myofibrillar due to their similar intensities (see Figure 2F). We input another set of manually drawn contours that demarcate the nucleus region and follow the same region-based thresholding approach to automatically identify the nucleus region. These pixels are then removed from the pixels that have been grouped as myofibrils, thus completing the segmentation process.

### Segmentation Accuracy Analysis

We measured the accuracy of the segmentation algorithm by comparing the automatically identified pixel regions of mitochondria, myofibrils and nucleus against those identified through manual segmentation.

Regions of mitochondria were manually demarcated within a small selection (randomly chosen) of slices within the 3D image stack. The pixels in these regions were then classified as mitochondrial. This classification was then compared against the automatic classifications that the algorithm produced. Five comparison metrics were calculated:

- Sensitivity: also referred to as the true-positive rate is the fraction (percentage) of manually identified mitochondrial pixels that were classified by the algorithm as mitochondrial.
- Specificity: also referred to as the true-negative rate is the fraction (percentage) of the manually identified non-mitochondrial pixels that were correctly classified by the algorithm as non-mitochondrial.
- Accuracy: is the percentage of all (mitochondrial and non-mitochondrial) automatically classified pixels that match the manual classifications.
- F1-score: is another metric like sensitivity. A value of 1 means that the method has a 100% true positive rate and a value of 0 means that the method does not perform correct classifications at all.

Using these metrics, we compared the performance of the algorithm in correctly classifying manually segmented regions for Cell 1 and Cell 2 in Tissue Block 1 (see Figure 1) relative to three other algorithms: (i) Fiji’s machine-learning based WEKA segmentation tool kit^6^; (ii) a variation of an automatic thresholding technique, Fiji Robust threshold segmentation; and (iii) IMOD’s imodauto command^7^. We also investigated the minimum number of slices in which the cell boundary must be manually segmented for accurate segmentations.

### 3D Quantification of Organelle Distribution

We measured the spatial variation in mitochondrial and nucleus area fractions through the depth of the cell by quantifying the number of pixels of each classification in each longitudinal section from the cell surface (see Figure 9). Specifically, the proportion of mitochondria or nucleus pixels relative to the total number of pixels in a longitudinal section was calculated.

In addition to the total area fractions, a measure of local density of mitochondria within each longitudinal section was calculated as the proportion of mitochondria pixels within a small square kernel window (3×3 micrometer in size) that was placed over the segmented image. This was repeated over the entire image and the range, first and second quartiles and median local mitochondrial density were calculated for each longitudinal section.

## RESULTS AND DISCUSSION

Figures 2D, 2E, 2F and Figure 3 shows the results of the segmentation algorithm on the two 3D SBF-SEM image datasets that were acquired at the University of Melbourne. Figure 3A-B shows the segmentation of Cell 1 from Figure 1 in detail. Figure 3C shows a 3D rendering of Cell 2 from Figure 1 and Figure 3D shows a rendering of Cell 3 from Figure 1.

**Figure 3:**
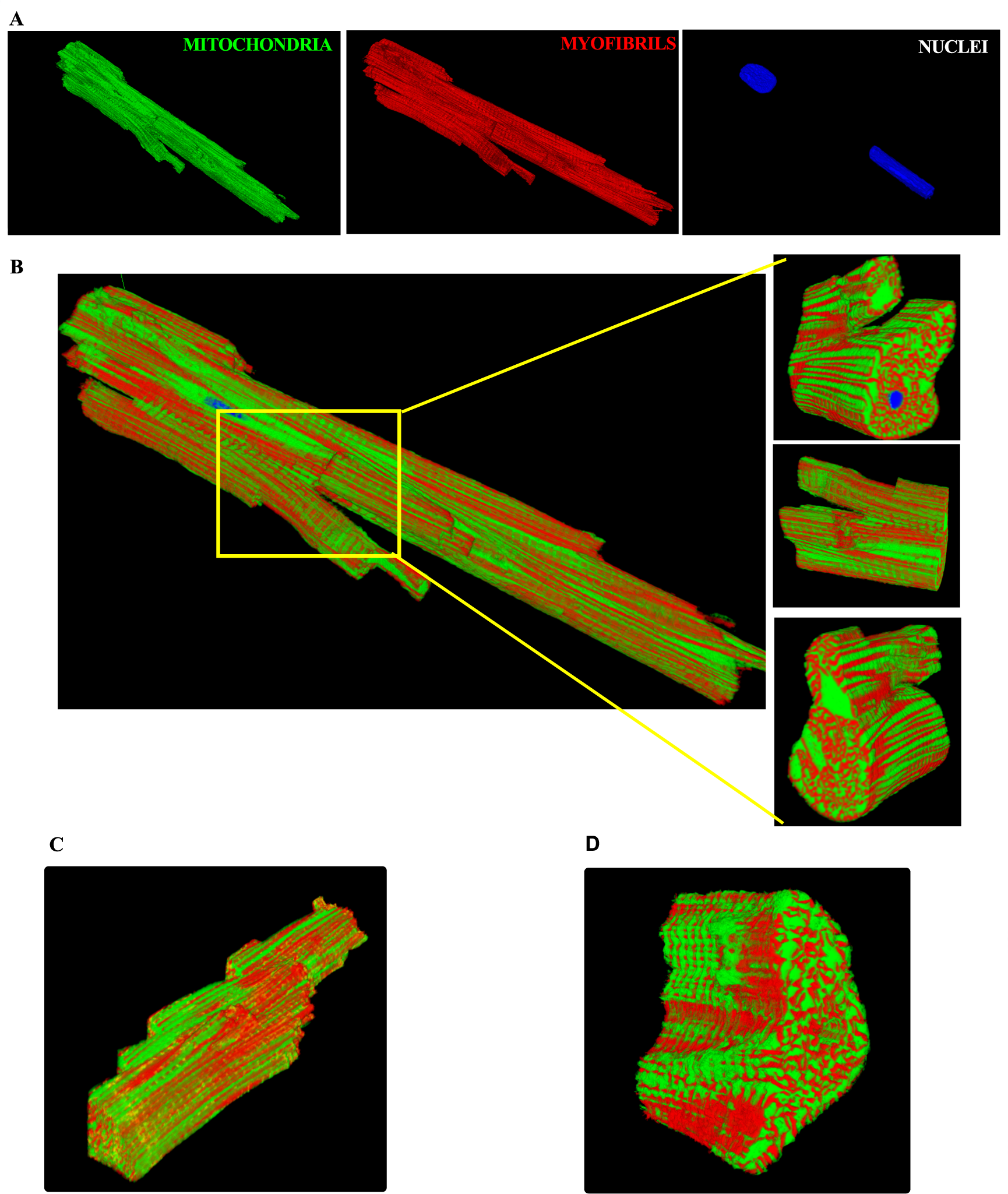
(A and B): 3D Rendering of myofibrils, mitochondria and nuclei within Cell 1 in the SBF-SEM dataset. The inset views provide a closer look at the organization of these components near a branching point. (C and D) are 3D renders of the segmentations of Cell 2 and Cell 3

### Quantitative Measurement of Accuracy and Performance Relative to Existing Tools

The accuracy of the new algorithm in classifying each pixel in Cell 1 as mitochondrial or myofibrillar when compared to manual segmentations is given in Table 1. The algorithm was also compared against the accuracy of three algorithms (relative to manual segmentations) available in Fiji and IMOD. As shown, the algorithm is highly accurate with a 97.6% accuracy rate when a manually drawn contour of the cell boundary is given every 10^th^ slice. The new algorithm also has the highest F1-score, which is a measure of the true positive rate. WEKA and robust segmentation achieved lower F1-scores. This may be a reflection of sub-optimal settings in these algorithms but it highlights the challenge for users to optimize a segmentation tool’s settings to extract regions of interest.

**Table 1:** Comparison of the performance of the new algorithm and other existing methods in correctly classifying manually segmented mitochondria in Cell 1.

| **Cell 1** | **F1-score** | **accuracy** | **sensitivity** | **specificity** |
| --- | --- | --- | --- | --- |
| **WEKA** | 0.709 | 96.6% | 69.2% | 98.3% |
| **Robust** | 0.697 | 96.0% | 75.6% | 97.3% |
| **IMOD** | 0.820 | 97.5% | 94.8% | 97.6% |
| **New algorithm** | 0.832 | 97.6% | 95.5% | 97.8% |

The performance of IMOD is similar to the new algorithm, which is expected because of the principles of our algorithm and the IMOD algorithm being similar. However, a closer, slice by slice comparison of the segmentation results for Cell 1 between IMOD and the new algorithm showed that IMOD incorrectly classified myofibril regions as mitochondrial, see Figure 4.

**Figure 4:**
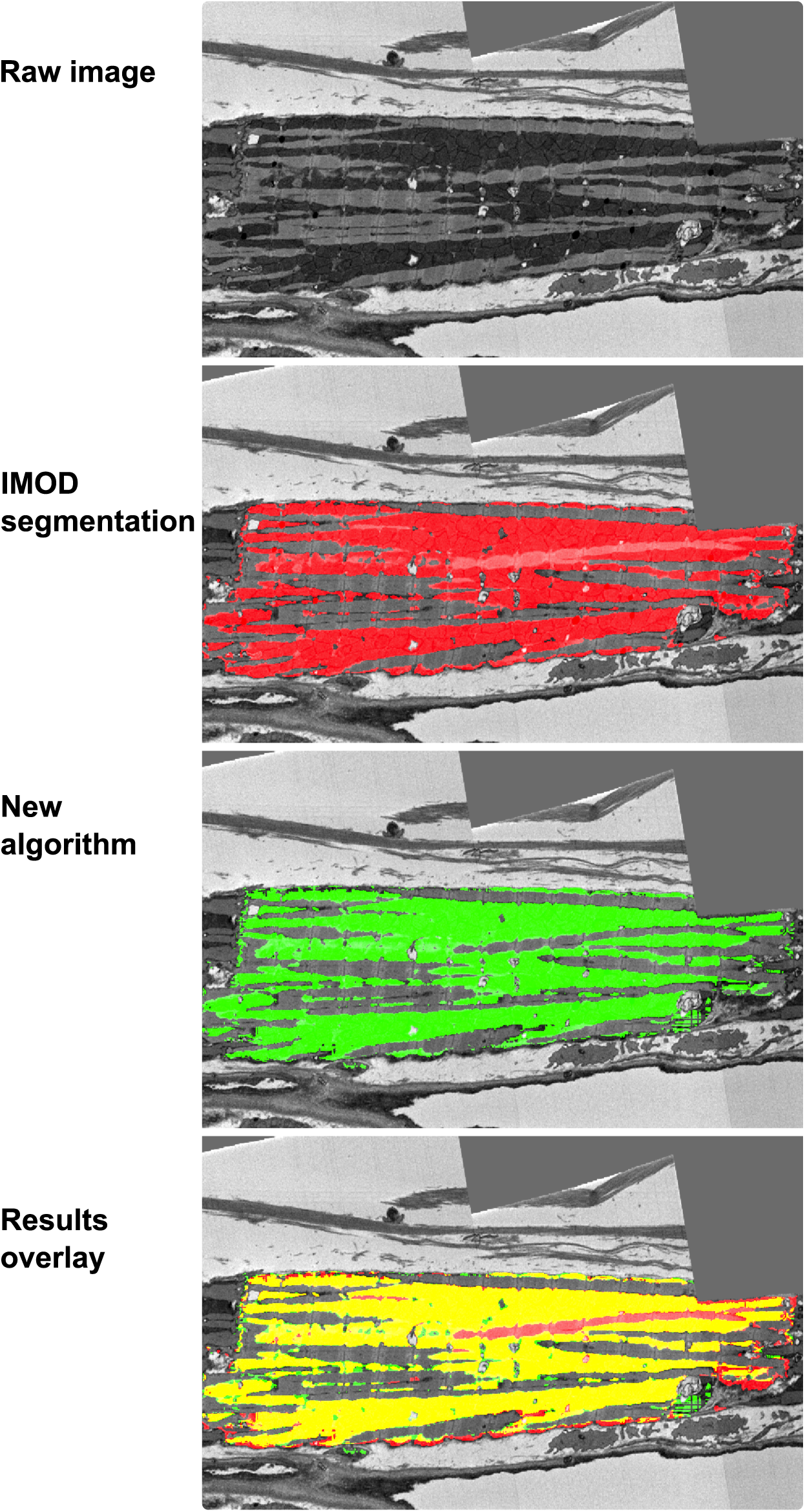
Comparison of segmentations of mitochondria by IMOD and our new algorithm. The first three images show the raw, IMOD segmentations and new algorithm segmentations of mitochondria. The results overlay shows a strip of red that highlights a myofibril region that has been classified by IMOD as mitochondrial. AH other regions are yellow because they have been correctly identified as mitochondrial m both IMOD and our new algorithm.

Segmenting every 10^th^ image slice in an image stack of one thousand and nineteen slices is a very tedious task. Table 2 shows the change in segmentation accuracy of the new algorithm if we chose to use manually drawn cell boundary contours every 20^th^ or every 50^th^ slice. With each slice having a z-resolution of 50 nm the accuracy is not severely affected even when using manually drawn contours only every 50^th^ slice. IMOD’s imodauto program differs from our algorithm in this regard. For any given slice, IMOD copies the nearest contour, the cell boundary, above or below the current slice and does not perform any extrapolation of the contour shape to capture the true shape of the cell boundary on the current slice. Therefore, IMOD does not segment entire regions of the cell when only every 50^th^ slice of the tissue block is manually segmented for the cell boundary. The new algorithm has been developed to enable rapid processing of SBF-SEM cardiac cell data.

**Table 2:** The accuracy of the algorithm in identifying manually segmented mitochondria regions in Cell 1 with fewer prior segmentations of the cell boundary.

| Cell 1 | F1-score | accuracy | sensitivity | specificity |
| --- | --- | --- | --- | --- |
| 10-slice | 0.832 | 97.6% | 95.5% | 97.8% |
| 20-slice | 0.854 | 92.4% | 97.7% | 90.9% |
| 50-slice | 0.828 | 90.8% | 98.1% | 88.7% |

### Capturing the Intercalated Discs

Figure 5 illustrates the accuracy of our algorithm in identifying the intercalated disc boundary. The left image shows a small portion of the 3D data with an intercalated disc and a portion of the manually demarcated cell boundary in green. Notice that the manual contour does not truthfully follow the cell boundary. The image on the right in Figure 5 shows the segmentation result along with the manually demarcated cell boundary (now in magenta). Comparison of the two images side by side shows that the algorithm only classifies pixels that fall within the true intercalated disc as opposed to including all pixels that fall within the manually demarcated boundary.

**Figure 5:**
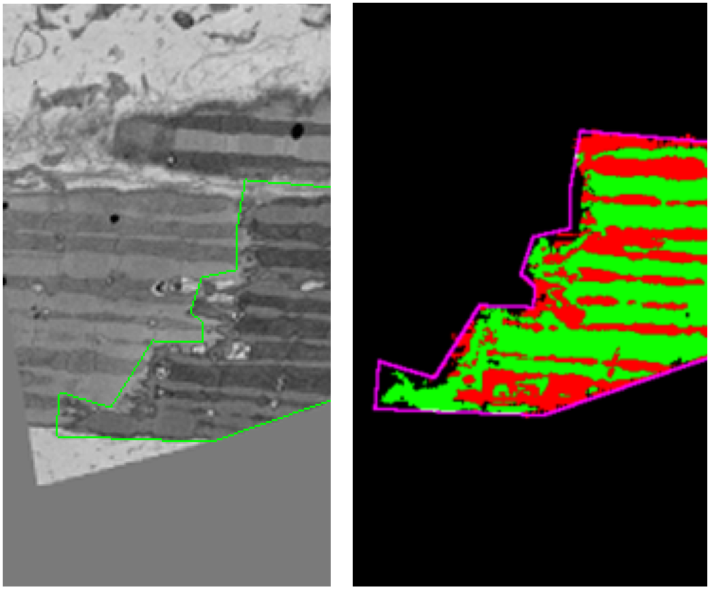
Accuracy in following the intercalated disc. Left panel shows the rough outline of the intercalated disc on the raw image. Right panel shows the segmented image, only containing pixels within the intercalated disc.

### Sensitivity to Image Contrast

The accuracy of the segmentation algorithm can be severely affected by differences in contrast between different image datasets. A closer examination of the raw data from Tissue Block 1 in Figure 6 shows that the intensity and contrast distribution changes considerably from cell to cell and even within the same cell. This is reflective of the cell to cell variation in heavy metal staining during tissue preparation. As depicted in Figure 6, Cell 2 exhibited several regionally different intensity profiles and is, therefore, a more challenging dataset to segment than a cell with a homogeneous intensity profile. Table 3 shows that our algorithm performs better than IMOD in this case as well.

**Table 3:** Comparison of the performance of the new algorithm against IMOD in correctly classifying manually segmented mitochondria in Cell 2.

| Cell 2 | F1-score | accuracy | sensitivity | specificity |
| --- | --- | --- | --- | --- |
| IMOD | 0.612 | 96.7% | 91.3% | 96.8% |
| New algorithm | 0.741 | 98.2% | 87.9% | 98.5% |

**Figure 6.**
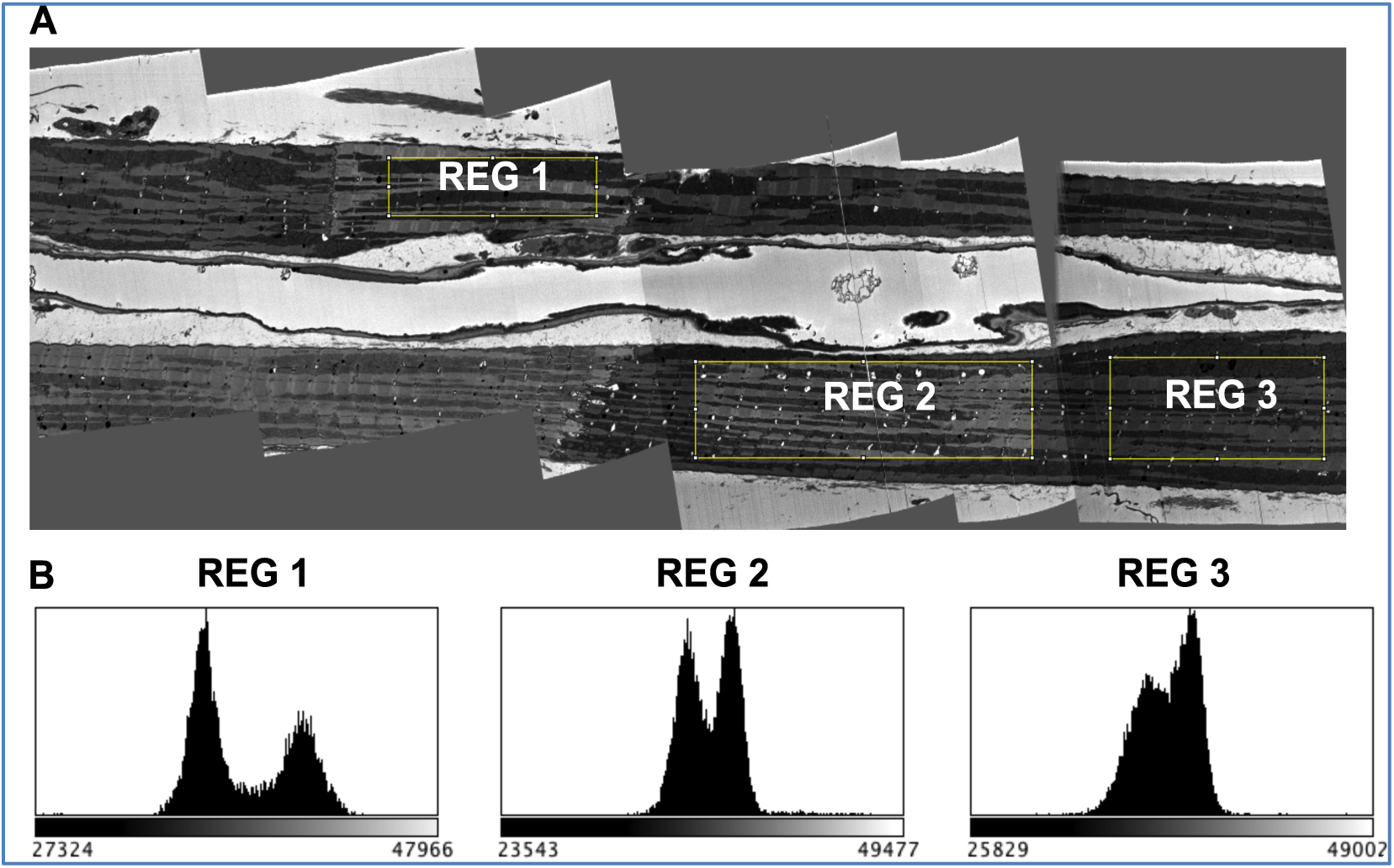
Intensity profiles vary within a SBF-SEM image slice. (A) shows an image slice from Tissue Block 1 (of Figure 1). The yellow boxes demarcate three regions, for which the corresponding intensity profiles are plotted in

Finally, we also tested whether our segmentation algorithm only works with datasets acquired using the University of Melbourne Teneo Volumescope machine. Figure 7 shows a successful segmentation of mitochondria from SBF-SEM data collected at UCL. Figure 8 shows the results of using the proposed algorithm on another, published dataset^9^. The key difference between this published dataset and all the other data in this study is the lack of two distinct peaks in the intensity histogram (compare Figure 8B with Figure 7B and Figure 6B). As our algorithm is based on the identification of two peaks (see Figure 2C), the algorithm mistakenly classifies myofibril regions as mitochondria in this dataset (Figure 8C). However, Figure 8C also qualitatively compares our algorithm against IMOD and Fiji’s robust segmentation algorithm and shows that our method is more robust to variations in contrast than other existing threshold-based algorithms.

**Figure 7:**
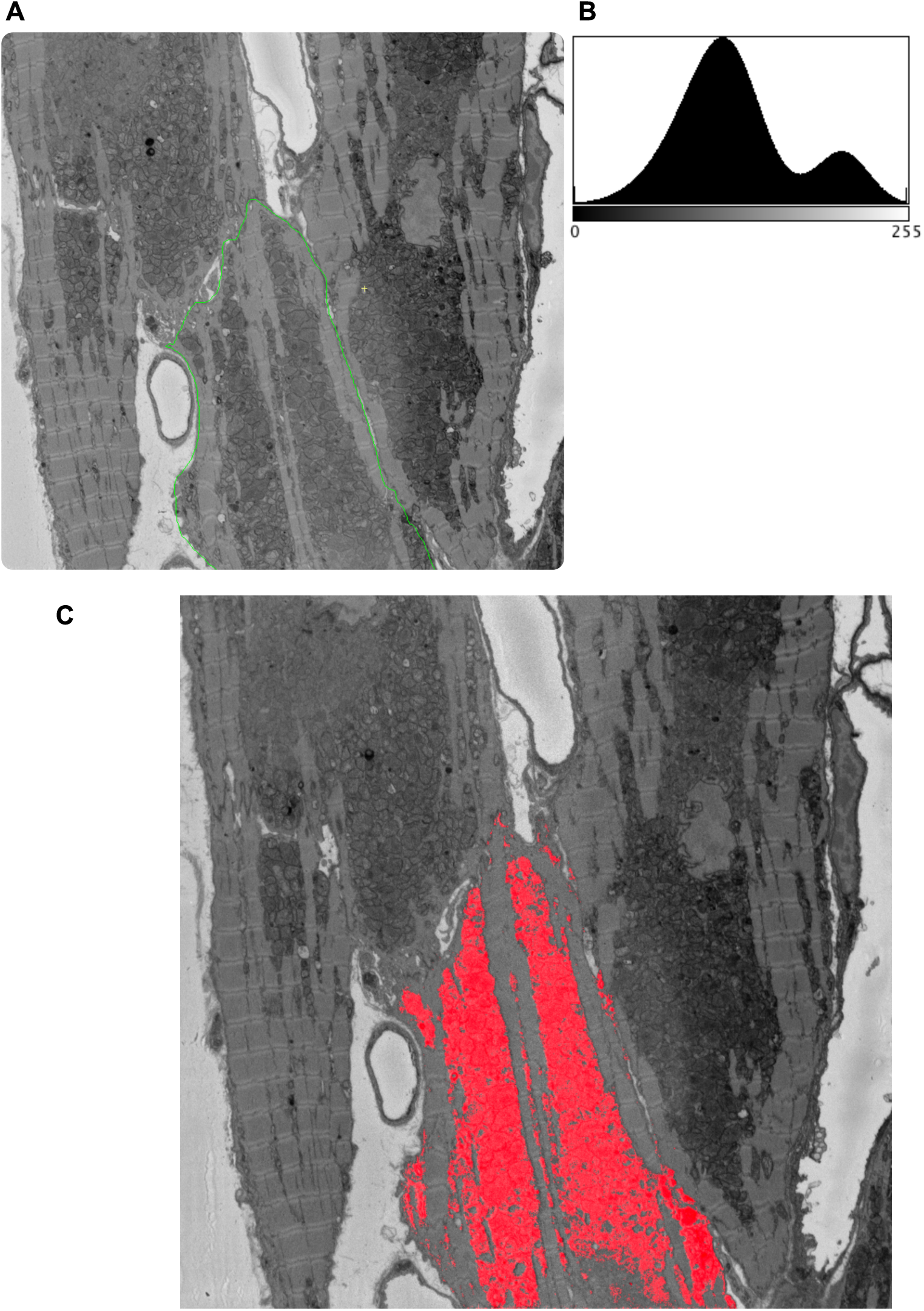
The new algorithm can segment mitochondria in SBF-SEM data collected at UCL. (A) shows the image with the manually demarcated boundary of one of the cells; (B) shows the 3D intensity profile across this dataset; (C) shows that the new algorithm can identify the mitochondrial region within the manually demarcated adequately.

**Figure 8:**
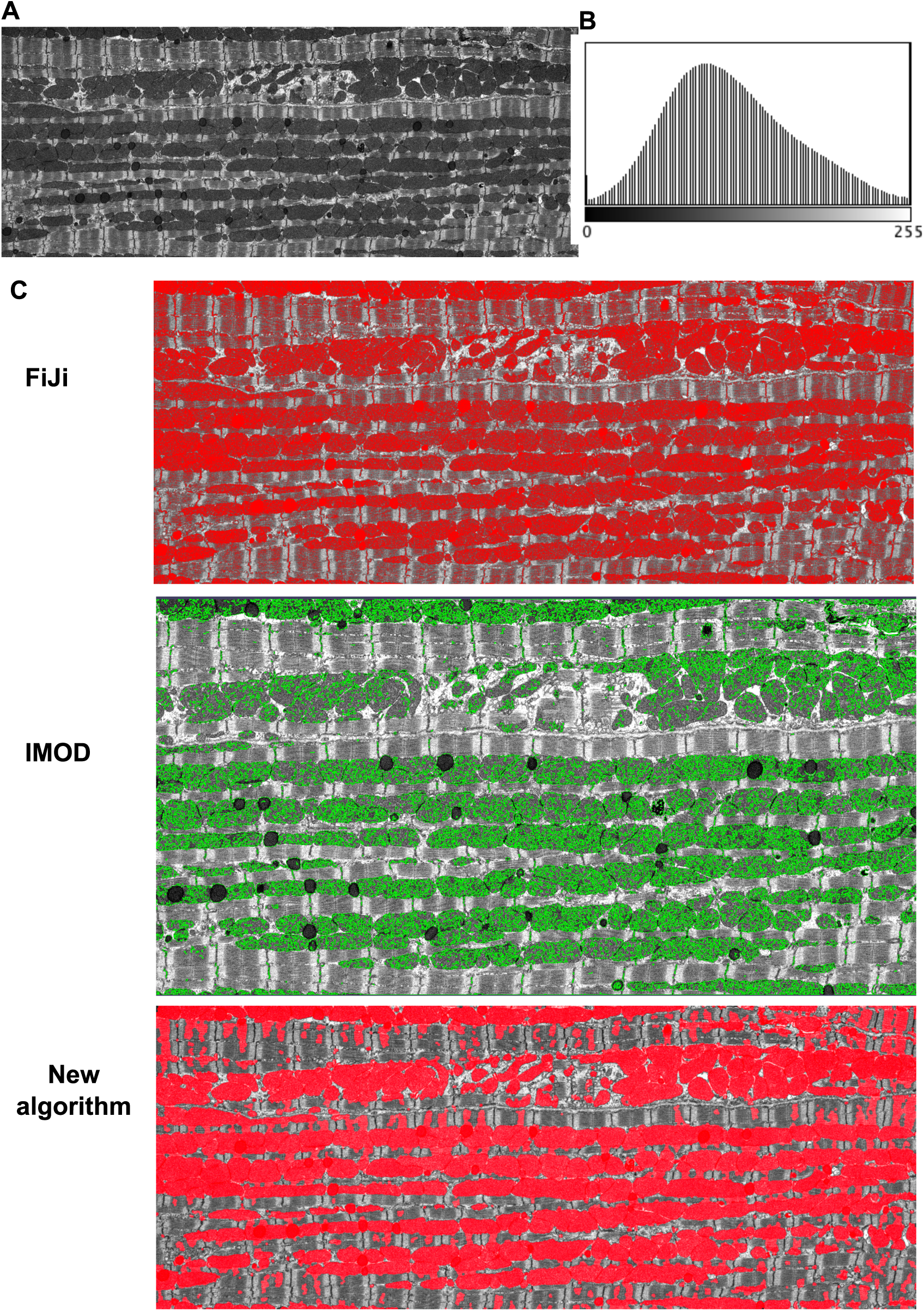
Segmentation of a dataset with no distinct peaks. (A) shows the raw image and (B) shows the intensity histogram. (C) shows a visual comparison of two threshold-based algorithms against the proposed algorithm of this paper, which is also threshold-based. The new algorithm is able to capture mitochondria more accurately than the other two methods.

### Quantification of Distribution of Organelles in 3D

With the algorithm performance assessed, we set out to extract information about the 3D distribution of the myofibrils and mitochondria within the segmented cells. Figures 3A and 3B show that the cell contractile proteins branch off as they contact a neighboring cell (not shown). We found a moderate increase in clustering of mitochondria around the nuclear volume and the cellular branching points. This is depicted by relatively larger crosssectional regions in green in the inset images in Figure 3B.

Table 4 documents the volume fractions of mitochondria, myofibrils and nuclei in the three cells.

**Table 4:**
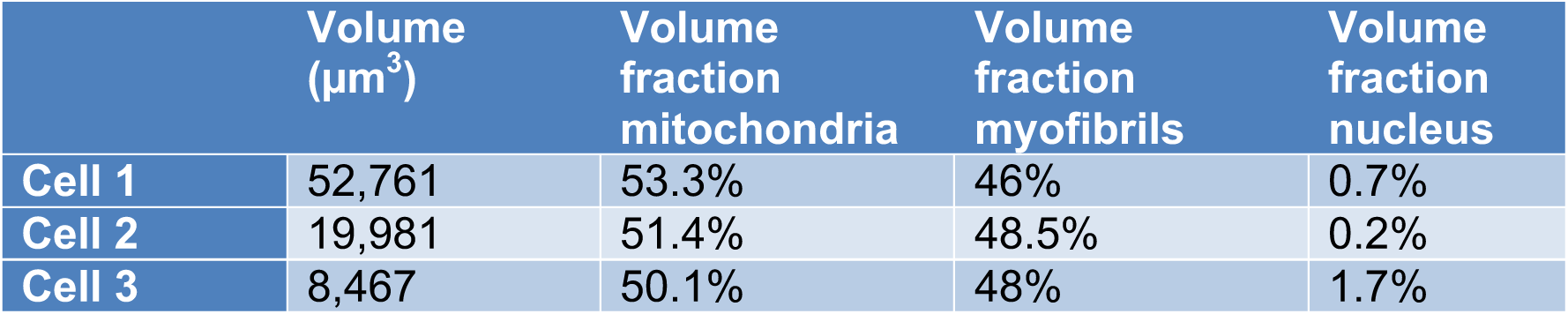
3D quantification of cell volume and volume fractions of mitochondria, myofibrils and nuclei.

Figure 9 shows the variation in area fraction of mitochondria and nuclei through the depth of the cell. The plot for Cell 1 shows that the mitochondria make up a large proportion of the cell at the cell periphery. The graph also shows that the area fraction of mitochondria is lowest, approximately 50% of cell volume, towards the core of the cell volume. This is not evident in Cell 2 because the dataset for tissue block 1 only captured a portion of the cell cross-section (see Figure 9). The dataset for Cell 3 does capture the entire cell cross-section and therefore depicts a similar variation in mitochondrial area fraction through the cell volume as Cell 1 (see Figure 9, Cell 3). The mitochondria underneath the membrane are referred to as sub-sarcolemmal (SSM) mitochondria, which have distinct molecular characteristics when compared to mitochondria that are deep within the cell, which are called inter-myofibrillar (IMF) mitochondria^14,15^. Our segmentations reflect the possibility that there is an abundance of SSM mitochondria forming a covering around the bulk of the contractile proteins and IMF-mitochondria.

**Figure 9:**
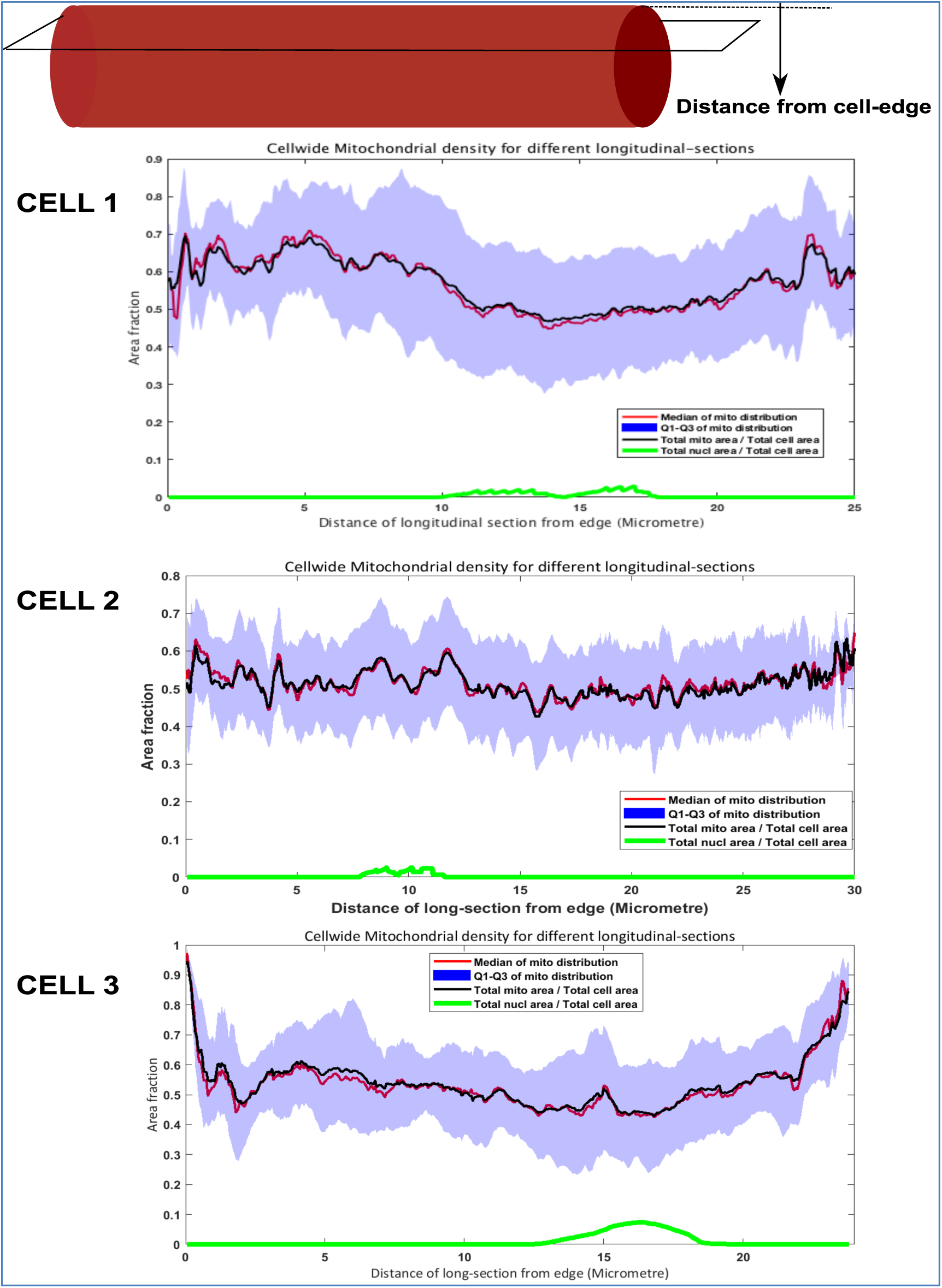
Spatial variation in volume fraction of mitochondria and nuclei within the 3 cells. The top panel shows an illustration of the direction along which the variation in mitochondrial and nucleus volume fraction w/as measured in the data.

Figure 9 also shows the variability in local density/distribution of mitochondria within each longitudinal section. This shows that the distribution of mitochondria within a section is non-uniform. Local clustering of mitochondria within the cell cross-section, even away from the nucleus region, is responsible for these local differences as reported previously^16,17^.

Our quantifications suggest that there are more mitochondria than myofibrils, which is in direct opposition to many previous studies that have reported volume fractions of the order of 50% myofibrils and 30% mitochondria^18,19^. With only three cells segmented, our results are not conclusive. However, all previous measurements were made using 2D data, which would not have captured the increased clustering of mitochondria in the nucleus region or the sub-sarcolemma. To test this hypothesis, we also performed an area fraction calculation on a small 2D section of Cell 1 (see Figure 10), in which myofibrils covered 59.5% of the area and mitochondria only occupied 38.6% of the area.

**Figure 10:**
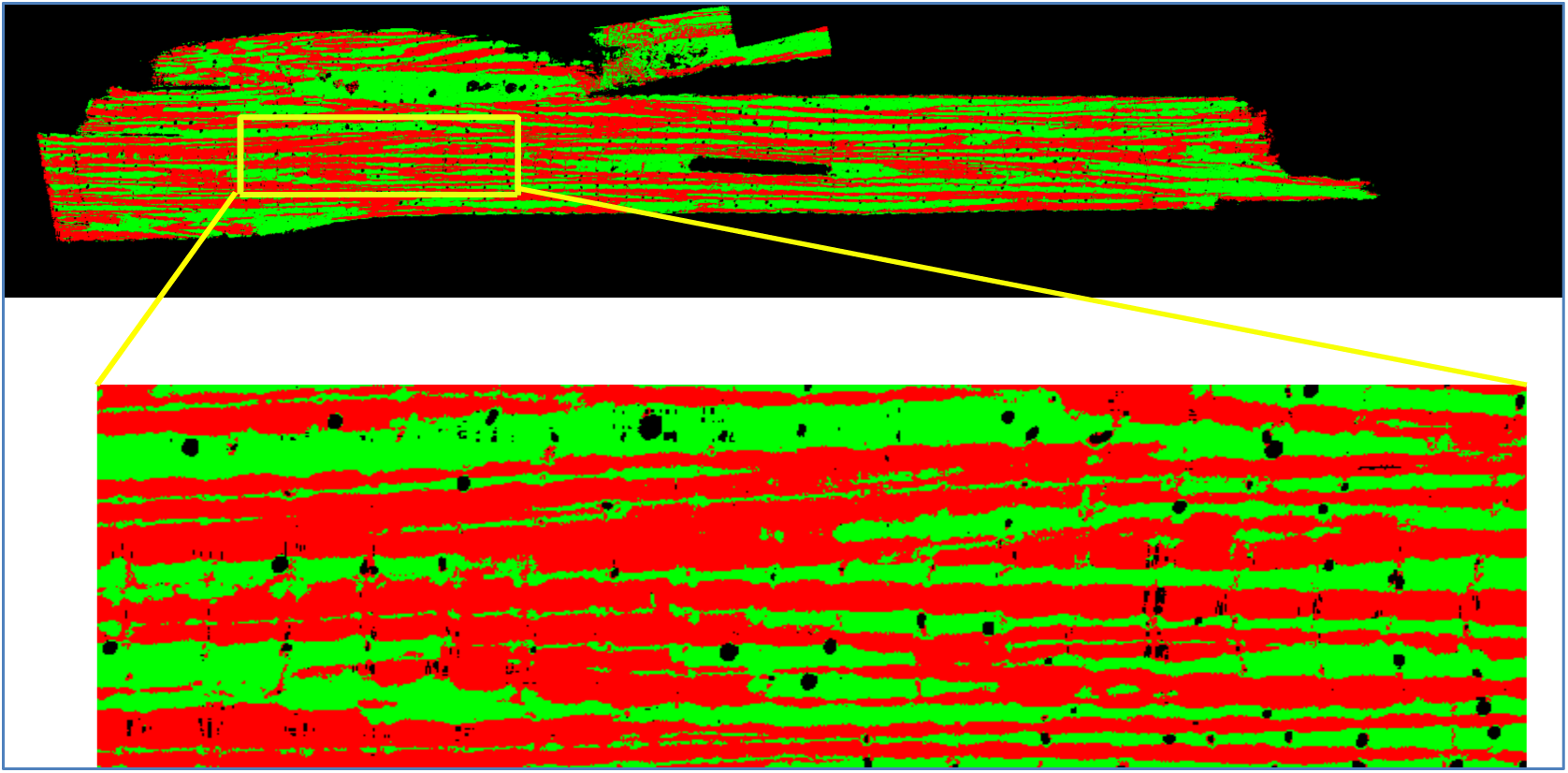
2D sample image used for 2D stereology. A small rectangular section (inside yellow box and zoomed in view in the bottom image) was extracted from the 3D segmentation of Cell 1 to calculate the area fractions of mitochondria and myofibrils.

### Creating Meshes from Segmentations for Computational Physiology Modelling

Finally, we demonstrate the feasibility of converting the segmentations from our algorithm into 3D finite element meshes in Figure 11. Segmentations for Cell 3 were used as input in a 3D tetrahedral mesh generation program called iso2mesh^20^. Such meshes can be used for simulations of intracellular calcium dynamics^10^ and bioenergetics^21^ even without other details such as t-tubules or the sarcoplasmic reticulum.

**Figure 11:**
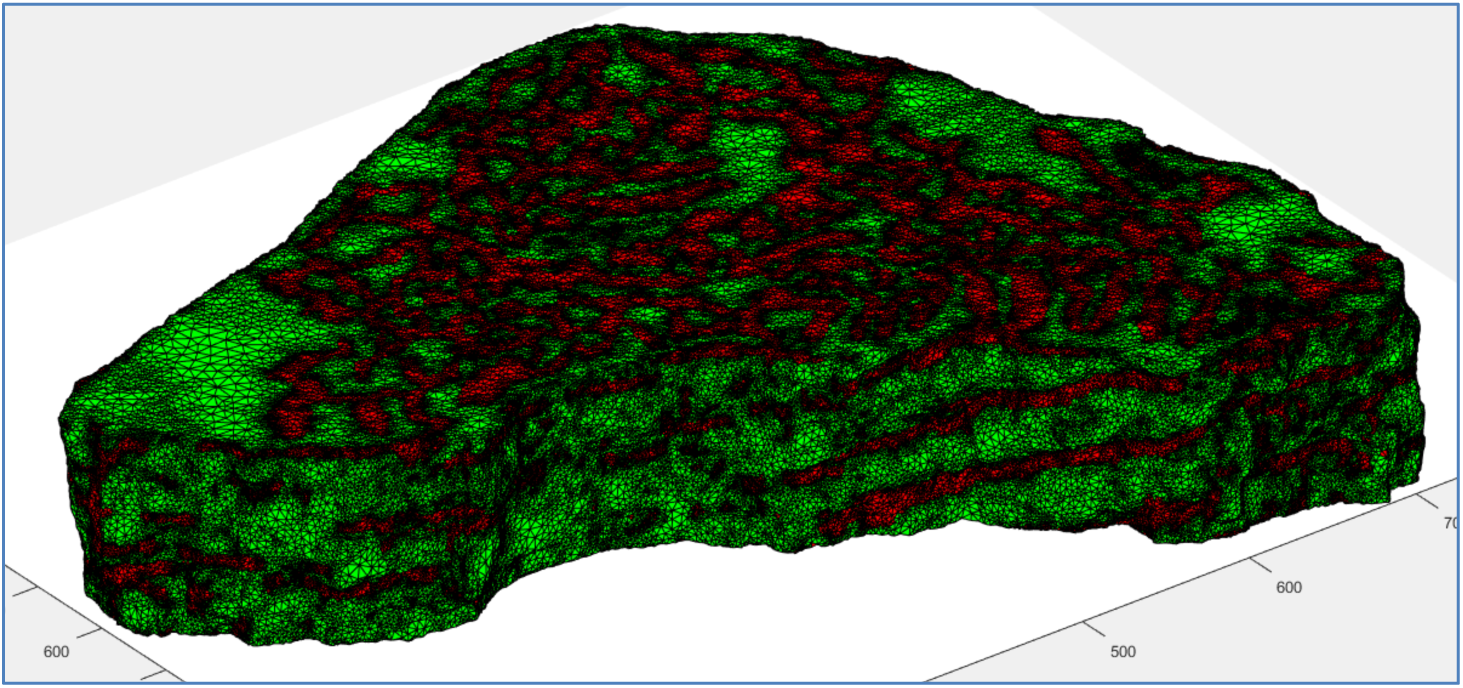
A 3D Finite element mesh of Cell 3 using the segmentations from the new algorithm. Green regions represent mitochondria and red regions represent myofibril regions

## CONCLUSIONS

We have developed an easy-to-use algorithm that only requires a stack of relatively approximate manual demarcations of the cell boundary as input to accurately classify the cell pixels into myofibrils, mitochondria or nuclei. Our analysis of the minimal number of manual segmentations required shows that the algorithm works well even when only every 50^th^ slice (slices at z-depth resolution of 50 nm) is demarcated.

We anticipate this algorithm to be of use to cardiac computational and experimental physiologists alike as these datasets provide a more comprehensive view of the cell than had been possible before the advent of SBF-SEM. Therefore, we have provided all our segmented data and MATLAB implementation of the algorithm for the research community at www.github.com/CellSMB/sbfsem-cardiac-cell-segmenter/.

## ACKNOWLEDGEMENTS

The authors would like to acknowledge the support of the Royal Society of New Zealand Marsden Fast Start grant 11-UOA-186, Collier Charitable Fund and the Australian Research Council Discovery Project grant DP170101358. The authors acknowledge the support from the Trace Analysis for Chemical, Earth and Environmental Sciences (TrACEES) platform from the Melbourne Collaborative Infrastructure Research Program at the University of Melbourne and thank Dr Jay Black (School of Earth Sciences) for operating the micro-CT scanner and processing data. We would like to thank Dr Peter Munro from the UCL Institute of Ophthalmology for his help with tissue processing and imaging at University College London. We also thank Dr. Brian Glancy at NIH/NHLBI for valuable discussions and access to the data in Figure 7. The FEI Teneo Volumescope was purchase thru the Australian Research Council Linkage Infrastructure, Equipment and Facility Scheme LE150100004.

